# One-step Isolation of Protein C-terminal Peptides from V8 Protease-digested Proteins by Metal Oxide-based Ligand Exchange Chromatography

**DOI:** 10.1101/2021.08.29.456766

**Authors:** Hiroshi Nishida, Yasushi Ishihama

## Abstract

We have developed a one-step method to isolate protein C-terminal peptides from V8 protease-digested proteins by metal oxide-based ligand-exchange (MOLEX) chromatography. V8 protease cleaves the C-terminal side of Asp and Glu, affording a digested peptide with two carboxy groups at the C-terminus, whereas the protein C-terminal peptide has only one α-carboxy group. In MOLEX chromatography, a stable chelate is formed between dicarboxylates and metal atoms, so that the non-terminal (i.e., internal) peptide is retained, whereas the protein C-terminal peptide flows through the MOLEX column. After optimization of the MOLEX chromatographic conditions, 1619 protein C-termini were identified from 30 μg of peptides (10 μg each, in triplicate) derived from human HeLa cells by means of nanoLC/MS/MS. When the MOLEX-isolated sample from 200 µg of HeLa peptides was further divided into six fractions by high-pH reversed-phase LC prior to nanoLC/MS/MS, 2202 protein C-termini were identified with less than 3% contamination with internal peptides. We believe this is the largest coverage with the highest purity reported to date in human protein C-terminomics. This fast, simple, sensitive and selective method to isolate protein C-terminal peptides should be useful for profiling protein C-termini on a proteome-wide scale.

## INTRODUCTION

Protein terminal sequences play an important role in biological phenomena such as translational initiation, stability, complex formation and signal transduction.^1–6^ Moreover, genetic mutation, RNA splicing, post-translational modification and endogenous proteolysis enable the generation of multiple proteoforms from a single gene.^7,8^ Proteolytic processing is especially related to the diversity of terminal sequences and functions of proteins. For instance, membrane-bound proteases cleave the same membrane proteins at more than one site, and the generated proteoforms with different neo-terminal sequences exhibit distinct physiological properties.^9,10^ Therefore, large-scale analysis of protein termini is important to characterize protein functions.

Shotgun proteomics based on liquid chromatography/tandem mass spectrometry (LC/MS/MS) is a powerful tool to acquire comprehensive information on the proteome. In global proteome samples, however, identification of protein terminal peptides is hampered by the presence of large amounts of internal peptides. Thus, several methods for the isolation of protein N- and C-terminal peptides have been developed. Among them, COFRADIC (combined fractional diagonal chromatography)^11^ and TAILS (terminal amine isotope labeling of substrates)^12^ have been widely used for N-terminomics, and the obtained N-terminomic data have been utilized for the analysis of degradomes and proteoforms.^13^ On the other hand, protein C-terminomics is still challenging, since selective modification of protein C-termini is difficult and the ionization efficiency of protein C-terminal peptides is generally low^.14^ To date, several methods have been developed to isolate protein C-terminal peptides from digested protein mixtures.^15–28^ These methods can be roughly divided into two types, involving positive and negative selection. Positive selection targets protein C-terminal carboxy groups for labeling with oxazolone or carboxypeptidase^16,17^, although the labeling efficiency of these methods is incomplete or biased by the nature of the amino acid residues at the protein C-termini. Negative selection is based on depletion of internal peptides followed by recovery of the remaining terminal peptides. Most methods for identifying protein C-terminal peptides employ negative selection. Van Damme et al. developed C-terminal COFRADIC^18^, which includes a strong cation exchange (SCX) chromatographic step, two RP-HPLC separation steps and two derivatization steps. Schilling et al. established a C-terminal version of TAILS, C-TAILS,^19^ in which protein C-terminal peptides are enriched via binding of internal tryptic peptides to polymers after chemical protection of the amino and carboxy groups of the proteins and digestion. Recently, modified versions of these methods with different types of labeling and different enzymes have been developed.^22,28^ In addition, an enrichment method of protein terminal peptides using only endopeptidase without chemical reaction has been developed,^25^ but this requires a reaction time of about a week. Dormeyer et al. developed an isolation method for protein C-terminal peptides together with acetylated protein N-terminal peptides using SCX chromatography.^15^ This method does not require a chemical reaction and is quick, but there is a bias because terminal peptides containing additional basic residues cannot be captured. Recently, a large-scale terminomics study using this method was published, but the efficiency of terminal peptide enrichment had to be sacrificed to reduce this bias, resulting in a purity of only 34% at best.^29^ In other words, previous methods all involve enrichment bias or labor-intensive procedures such as multiple steps of chemical or enzymatic reactions. Thus, there is still a need for a simple, easy and less biased method.

Recently, our group reported a new method referred to CHAMP (Chop and throw to AMPlify the terminal peptides) to isolate protein N-terminal peptides in only two steps, i.e., TrypN or lysarginase digestion and low-pH SCX chromatography,^30^ based on the concept that terminal peptides can be isolated by employing appropriate proteases followed by one-step chromatography. Following the success of this CHAMP N-terminal approach, we immediately tried a similar approach to isolate protein C-terminal peptides by combining V8 protease digestion with high-pH SAX chromatography, since these peptides appeared to be analogous to the protein N-terminal peptides except for having the opposite charge. However, this proved unsuccessful, because the pK_a_ of the guanidino group of Arg was too high to neutralize the charges.

In this study, we examined V8 protease and metal oxide-based ligand-exchange (MOLEX) chromatography for isolating protein C-terminal peptides based on the CHAMP concept. After optimizing the isolation conditions, we applied this method to HeLa lysates as a model for large-scale protein C-terminal profiling, and compared the results with those obtained by conventional methods.

## EXPERIMENTAL SECTION

### Materials

Cerium oxide (CeO_2_), gallium oxide (Ga_2_O_3_) and aluminium oxide (Al_2_O_3_) were purchased from FUJIFILM Wako (Osaka, Japan). Titansphere^®^ Phos-TiO (TiO_2_) was obtained from GL Sciences (Tokyo, Japan). ZirChrom^®^-PHASE (ZrO_2_) was purchased from ZirChrom Separations (Anoka, MN). UltraPure™ Tris Buffer was purchased from Thermo Fisher Scientific (Waltham, MA). Polyethylene frit was purchased from Agilent Technologies (Santa Clara, CA). SDB-XC Empore disks were from GL Sciences. Water was purified by a Millipore Milli-Q system (Bedford, MA). Protease inhibitors, phosphatase inhibitor cocktail 2 and phosphatase inhibitor cocktail 3 were purchased from Sigma-Aldrich (St. Louis, MO). All other chemicals including V8 protease were purchased from FUJIFILM Wako, unless otherwise specified.

### Cell Culture, Protein Extraction and V8 Protease Digestion

HeLa S3 cells obtained from the JCRB Cell Bank (Osaka, Japan) were cultured in Dulbecco’s modified Eagle’s medium with 10 % fetal bovine serum and 100 μg/mL Kanamycin at 37 °C, 5 % CO_2_. Proteins were extracted with phase-transfer surfactant (PTS protocol), as described previously,^31^ except that V8 protease was used instead of trypsin/LysC. Briefly, HeLa cells were suspended in PTS lysis buffer consisting of protease inhibitors, phosphatase inhibitor cocktail 2, phosphatase inhibitor cocktail 3, 12 mM SDC, 12 mM SLS in 100 mM Tris-HCl buffer (pH 9). The cell lysate was diluted 5-fold with 50 mM ammonium bicarbonate solution and incubated overnight with V8 protease (enzyme/substrate ratio = 1:20 (w/w)) at 37 °C. After digestion, an equal volume of ethyl acetate was added, and the solution was acidified with trifluoroacetic acid (TFA). Then the peptides were desalted with SDB-StageTips.^32,33^

### Metal Oxide-based Ligand-Exchange Chromatography

MOLEX chromatography was performed by using disposable pipette tip-based columns such as StageTips. MOLEX tips were prepared as follows. A polyethylene frit was stamped out using a blunt-end 16-gauge syringe needle and placed into a P200 pipette tip. Metal oxide particles were suspended in methanol, loaded into the pipette tip, and packed by centrifugation. Mobile phases were prepared with aqueous 40% acetonitrile in buffers of various pH values, adjusted with 1 M hydrochloric acid. Note that the pH was measured in the absence of acetonitrile. The MOLEX tips were equilibrated with 200 μL of the mobile phase. The desalted peptide samples were dried in a SpeedVac™ Concentrator (Thermo Fisher Scientific), reconstituted with the mobile phase at pH 5, and loaded onto the MOLEX tips. The eluted fractions were collected and diluted 10 times with 1 % TFA. The diluted fractions were desalted on SDB-StageTips. MOLEX tips packed with 10 mg TiO_2_, 10 mg ZrO_2_, 15 mg CeO_2_, 30 mg Al_2_O_3_ or 40 mg Ga_2_O_3_ were loaded with 5 μg of peptides.

### High pH Reversed Phase Fractionation

A MOLEX-isolated sample obtained from 200 µg of HeLa peptides was fractionated using an LC-Mikros™ pump (Shimadzu, Japan, Kyoto), an HTC-PAL autosampler with an off-line fraction collector unit and a Waters nanoEase™ M/Z Peptide BEH C18 analytical column (300 µm × 100 mm, 130 Å, 1.7 µm). Solvent C (2.5 mM ammonium bicarbonate) and solvent D (2.5 mM ammonium bicarbonate and 80% acetonitrile) were employed as the mobile phases of this system. A linear gradient was run from 2.5 to 45 % D in 30 min, and 45 to 99 % D in 1 min, followed by 99 % D for 4 min. The flow rate was set at 5 µL/min. The peptide sample reconstituted in 2.5 mM ammonium bicarbonate was fractionated at 90 sec intervals to obtain 24 fractions, which were combined to afford 6 fractions. After SpeedVac concentration, each fraction was reconstituted in 4 % acetonitrile with 0.5 % TFA and injected to a nanoLC/MS/MS system equipped with an Orbitrap Exploris 480 mass spectrometer (Thermo Fisher Scientific) as described below.

### NanoLC/MS/MS Analysis

The LC/MS/MS analyses were performed on an HTC-PAL autosampler (CTC Analytics, Zwingen, Switzerland) and an Ultimate 3000 pump (Thermo Fisher Scientific, Waltham, MA), combined with a TripleTOF 5600+ (SCIEX, Framingham, MA), a Q-Exactive, or the Orbitrap Exploris 480 mass spectrometer. The LC mobile phases consisted of solvent A (0.5% acetic acid) and solvent B (0.5% acetic acid and 80% acetonitrile). All mass spectrometric analyses were carried out in the data dependent acquisition (DDA) mode. In the experiments with the TripleTOF 5600+ system, peptides were separated on self-pulled needle columns (150 mm, 100 μm ID) packed with Reprosil-Pur 120 C18-AQ 3 μm (Dr. Maisch, Ammerbuch, Germany).^34^ The injection volume was 5 μL, and the flow rate was 500 nL/min. Analytes were reconstituted in 4 % acetonitrile with 0.5 % TFA. A linear gradient was employed from 5 to 40 % B in 20 min, and 40 to 99 % B in 1 min, followed by 99 % B for 4 min. The electrospray voltage was set to 2.4 kV in the positive mode. The mass range of full scan was from 300 to 1500 *m/z* and the mass range of MS/MS was from 80 to 1500 *m/z*. For the Q-Exactive system, LC conditions were the same as those for the TripleTOF 5600+ system. The electrospray voltage was set to 2.4 kV in the positive mode. The full MS scan was acquired with the mass range of 350-1500 *m/z*, resolution of 70000, AGC target of 3e^6^ and maximum injection time of 100 ms. MS/MS scan was performed by the Top10 method with the resolution of 17500, AGC target of 1e^5^, maximum injection time of 100 ms and isolation window of 2.0 Th. The precursor ions were fragmented by higher energy collisional dissociation with a normalized collision energy of 27 %. For the Orbitrap Exploris 480 system, self-pulled needle columns (250 mm, 100 μm ID) were packed with Reprosil-Pur 120 C18-AQ 1.9 μm. The injection volume was 5 μL, and the flow rate was 400 nL/min. The mobile phases were the same as those for the other systems. The flow gradient was set as follows: 5 % B in 8.3 min, 5-19 % B in 92.2 min, 19-29 % B in 34.5 min, 29-40 % B in 15 min, and 40-99 % B in 0.1 min, followed by 99 % B for 4.9 min. The electrospray voltage was set to 2.4 kV in the positive mode. The mass spectrometric analysis was carried out with the FAIMS Pro™ interface. FAIMS mode was set to standard resolution, and total carrier gas flow was 4.0 L/min. The compensation voltage (CV) was set to −40, −60 and −80, and the cycle time of each CV experiment was set to 1 s. The mass range of the survey scan was from 300 to 1500 *m/z* with resolution of 60000, 300 % normalized AGC target, and auto maximum injection time. The first mass of MS/MS scan was set to 120 *m/z* with resolution of 30000, standard automatic gain control, and auto maximum injection time. The fragmentation was performed by higher energy collisional dissociation with a normalized collision energy of 30 %. The dynamic exclusion time was set to 20 s.

### Data Analysis

The raw MS data files acquired by the TripleTOF 5600+ and Q-Exactive systems were converted into MGF (Mascot Genetic Format) files with msconvert in ProteoWizard. For peptide identification from the MS/MS spectra, an automatic database search using Mascot ver. 2.6 (Matrix Science, London, UK) was used against the UniProtKB/Swiss-Prot database of canonical isoforms (20,199 sequences). Mass tolerances were set as 20 ppm, 5 ppm for precursor ions and 0.1 Da, 20 ppm for fragment ions for the data acquired with the TripleTOF 5600+ and Q-Exactive systems, respectively. The cleavage site was set at the C-terminal side of Asp and Glu, allowing for up to 5 missed cleavages. Cys carbamidomethylation was set as a fixed modification. Met oxidation and protein N-terminal acetylation were set as variable modifications. The identification threshold was set to E-value < 0.05. The peak area values of identified peptides were obtained by the in-house match-between-run function, based on the targeting extracted ion chromatograms (XIC) approach.^35^ The raw MS data files acquired by the Orbitrap Exploris 480 mass spectrometer were searched by MSFragger^36,37^ (ver. 3.3) and Philosopher^38^ (ver. 4.0.0) and the UniProtKB database of human proteins including isoforms (42360 entries, 2021_08) was downloaded via FragPipe (ver 16.0). Search parameters were loaded with closed search default config, although enzyme “gluc cut after DE” was selected for protein digestion and only Met oxidation was selected as a variable modification. In advanced peak-matching options, the minimum number of fragments modeling was set to one, the minimum number of matched fragments was one, and the maximum number of fragment charge was set to five. The FDR filter was set to 0.01 at the protein level.

### Data Availability

The raw MS data and analysis files have been deposited with the ProteomeXchangeConsortium (http://proteomecentral.proteomexchange.org) via the jPOST partner repository (http://jpostdb.org)^39^ with the data set identifier PXD027960.

## RESULTS AND DISCUSSION

### V8 Protease Digestion and MOLEX Chromatography

The CHAMP isolation workflow for protein C-terminal peptides is shown in Figure 1A. Proteins extracted from samples were digested with V8 protease, which cleaves the C-terminal side of Asp and Glu residues of proteins. Individual internal peptides have two carboxy groups at their C-termini, whereas protein C-terminal peptides have only one carboxy group. Then, we used MOLEX chromatography to isolate the protein C-terminal peptides from internal peptides by trapping the internal peptides based on stable chelate formation between the metal atom and dicarboxylate. Protein C-terminal peptides were collected in the flow-through and early eluted fractions.

**Figure 1.**
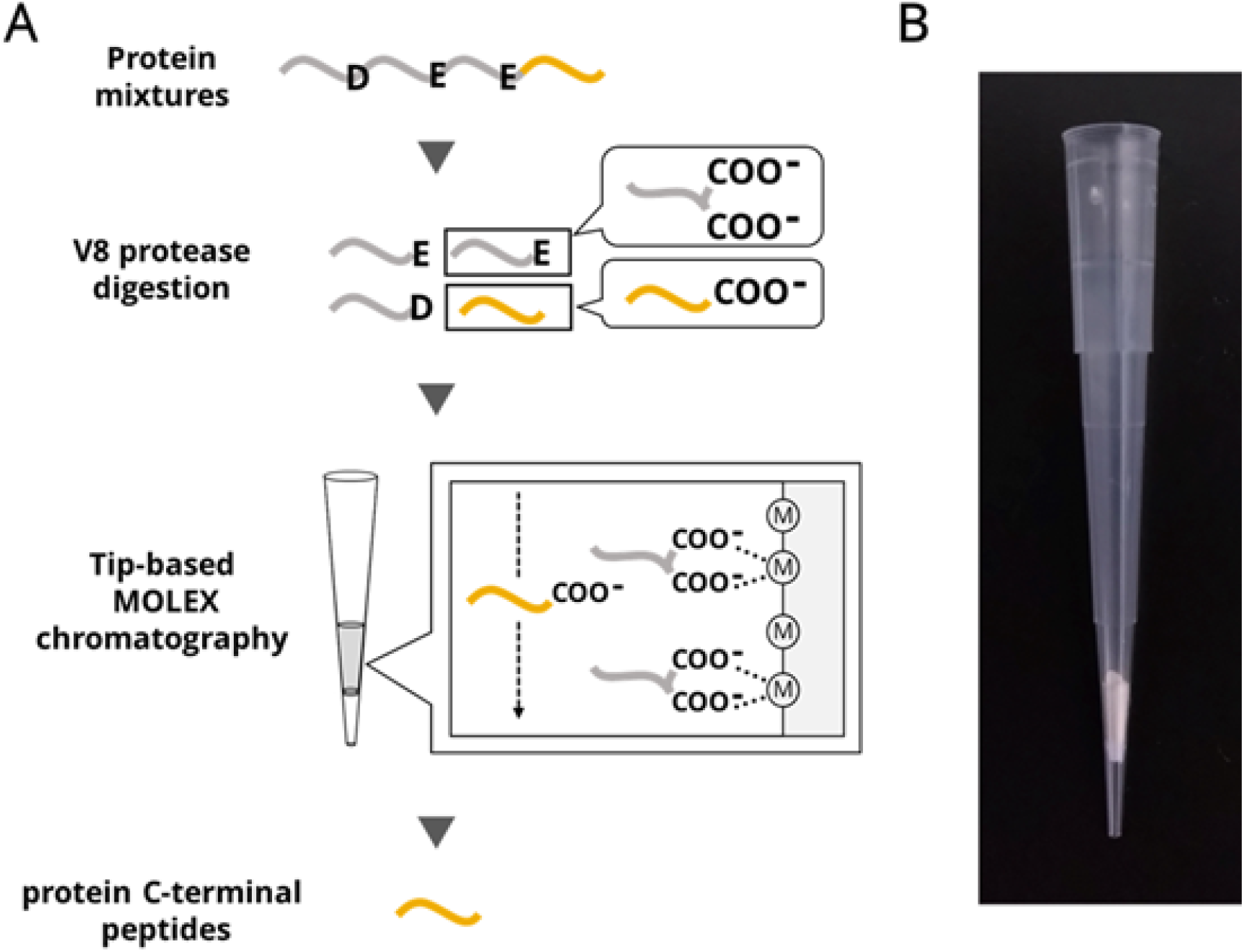
Schematic illustration of protein C-terminal peptide enrichment. (A) Proteins are digested with V8 protease and tip-based MOLEX chromatography is performed to trap the internal peptides. Circles with “M” represent metal atoms. The internal peptides form chelates with metal atoms, whereas the protein C-terminal peptides flow through the MOLEX column. (B) Photograph of a MOLEX tip column. For column preparation, a polyethylene frit cut out with a 16-gauge needle is inserted into a P200 pipette tip and then metal oxide slurry is packed by centrifugation.

MOLEX chromatographic procedures using TiO_2_, ZrO_2_, CeO_2_, Ga_2_O_3_ and Al_2_O_3_ (Ti-, Zr-, Ce-, Ga- and Al-MOLEX) were examined to isolate protein C-terminal peptides based on the formation of stable chelation and high chemical stability.^40–43^ StageTip-based MOLEX columns packed with the five different metal oxide particles were prepared as shown in Figure 1B. For the loading and elution buffers, aqueous 40% acetonitrile solutions in 30 mM sodium acetate at pH 2, 3, 4 and 5 were prepared, and the solution at pH 5 was used to solubilize 5 μg of the sample peptides. Four-step isocratic elution with pH 5, 4, 3 and 2, including the sample-loading step at pH 5, was conducted, and each pH fraction was collected, desalted and analyzed by nanoLC/MS/MS. Protein C-terminal peptides eluted earlier than internal peptides in all five MOLEX chromatographic runs, as shown in Figure 2A and Figure S1. These results confirmed that protein C-terminal peptides could be isolated by MOLEX chromatography, although the separation between protein C-terminal peptides and internal peptides was not sufficient. In Ti- and Zr-MOLEX chromatography, the protein C-terminal peptides were distributed into all fractions and internal peptides were observed only in the pH 2 fraction (Figure 2A). On the other hand, in Ce-, Ga- and Al-MOLEX chromatography, protein C-terminal peptides and internal peptides were eluted together with the elution buffers at pH 4 and less (Figure 2A and Figure S1). Based on these results, we selected Zr- and Ce-MOLEX for further optimization. Next, we examined the influence of amino acid composition of protein C-terminal peptides on MOLEX separation. We found that peptides with higher contents of Asp, Glu or Lys residues were present in the lower pH fractions (Figure 2B), suggesting that C-terminal peptides with Asp, Glu or Lys (DEK-Cterm peptides) have higher affinity for metal oxides and co-elute with internal peptides. We also employed immobilized metal ion affinity chromatography (IMAC), in which nitrilotriacetate or iminodiacetate resin was used to immobilize Fe^3+^, Ce^4+^ or Cu^2+^. As a result, protein C-terminal peptides were also successfully isolated by IMAC, although the separation between DEK-Cterm peptides and the internal peptides was incomplete, as in MOLEX chromatography with the five metal oxides. Considering the limited loading capacity of IMAC, we chose MOLEX chromatography rather than IMAC for further analyses.

**Figure 2.**
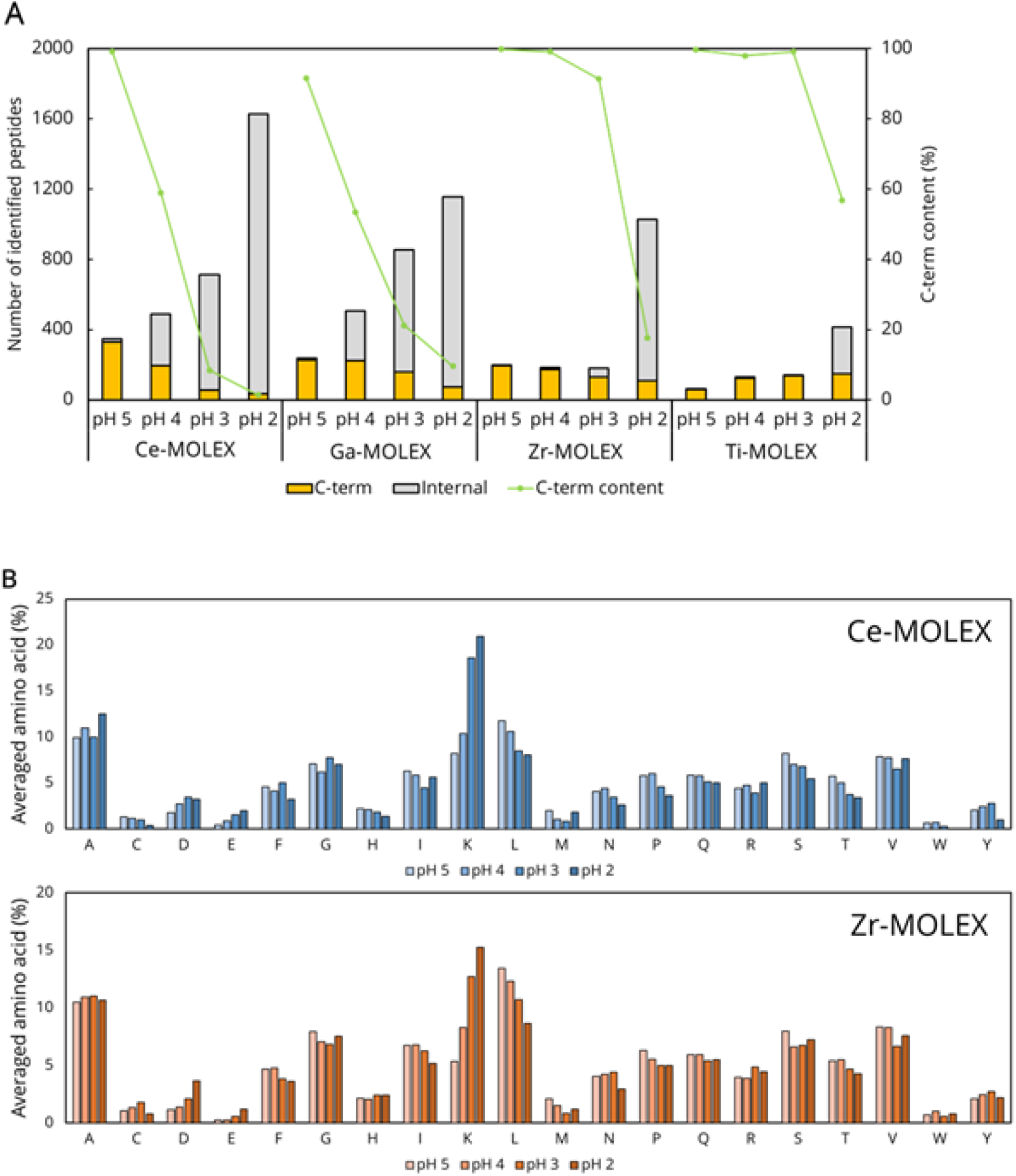
Isolation of protein C-terminal peptides from internal peptides by MOLEX chromatography using different metal oxides. (A) Numbers of identified protein C-terminal and internal peptides. Protein C-terminal peptide content (%) was calculated as the sum of peak area of all identified protein C-terminal peptides divided by that of all identified peptides. (B) Averaged amino acid (%) calculated from the number of each amino acid per peptide divided by the total number of residues for each pH fraction in Ce-MOLEX (top) or Zr-MOLEX (bottom) chromatography. The TripleTOF 5600+ system was used.

### Modifier Effects on MOLEX Chromatography

To improve the separation between protein C-terminal peptides and internal peptides, we added 600 mM sodium chloride (NaCl) to the elution buffers. We also prepared 200 mM ammonium dihydrogen phosphate solution as the final elution buffer after the pH-stepwise elution to elute all peptides by exchanging the ligands with phosphate.^44^ In Zr-and Ce-MOLEX chromatography, the elution conditions with and without NaCl were compared (Figure 3A). In both cases, the separation between protein C-terminal peptides and internal peptides was significantly improved in the presence of NaCl, and internal peptides did not elute until the final phosphate buffer. This result implies that the retention of the internal peptides was enhanced owing to the breakage of the intramolecular salt bridges. Furthermore, the average content of Lys per peptide became constant during the pH-stepwise elution in both cases of Zr- and Ce-MOLEX chromatography with 600 mM NaCl, suggesting that the lysine-metal interaction was suppressed (Figure 3B and S2). However, the protein C-terminal peptides containing Asp or Glu, i.e., acidic C-terminal peptides, were still co-eluted with internal peptides during the phosphate elution.

**Figure 3.**
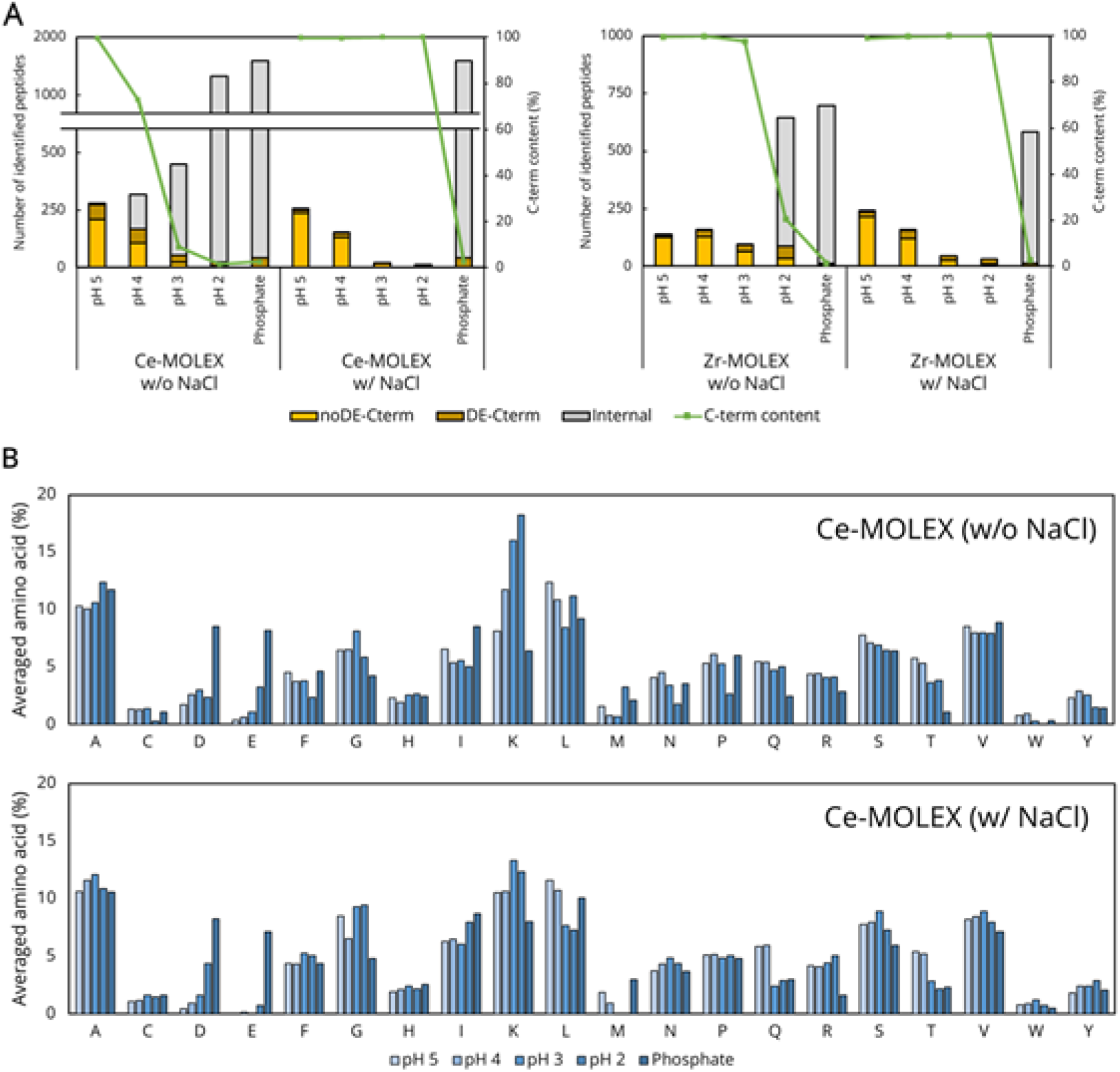
Effect of NaCl-containing mobile phase. (A) The number of identified peptides and the protein C-terminal peptide content in each fraction of Ce-MOLEX (left) or Zr-MOLEX (right) chromatography. V8-digested HeLa proteins (5 μg) were used for each condition. Protein C-terminal peptide content (%) was calculated as the sum of the peak areas of all identified protein C-terminal peptides divided by that of all identified peptides. (B) Averaged amino acid (%) was calculated from the number of each amino acid per peptide divided by the total number of residues for each fraction in Ce-MOLEX chromatography without NaCl (top) or with NaCl (bottom). The TripleTOF 5600+ system was used.

In hydroxy acid-modified metal oxide chromatography (HAMMOC) for phosphopeptide enrichment,^40^ competitive ligands such as lactic acid and β-hydroxypropanoic acid are effective as mobile phase modifiers to suppress the retention of acidic non-phosphorylated peptides. Therefore, we examined the effect of adding five amino acids with different lengths between the intramolecular amino and carboxy groups to the elution buffers as mobile phase modifiers (Figure 4A). We prepared aqueous 40% acetonitrile in 600 mM NaCl and 30 mM different amino acid buffers at pH 5 as the sample loading buffer for Ce- and Zr-MOLEX chromatography. Compared with the acetate buffer, the amino acid-modified buffers increased the number of identified acidic C-terminal peptides from the flow-through fraction while maintaining a high enrichment ratio except in the case of the glycine buffer (Figure 4B). We further investigated the influence of Asp or Glu position in the protein C-terminal peptides on the isolation efficiency. As shown in Figure S3, acidic protein C-terminal peptides with Asp or Glu at the first, second and third positions from the C-terminus could not be isolated under any of the conditions, representing a limitation of this approach. Overall, the largest identification number of acidic protein C-terminal peptides was obtained in Ce-MOLEX with δ-aminovaleric acid (Ce-MOLEX_δ-AVA) and Zr-MOLEX with γ-aminobutyric acid (Zr-MOLEX_γ-ABA), respectively, indicating that each metal oxide has a specific preference for the modifiers, which worked as competitors of the acidic C-terminal peptides. We further compared Ce-MOLEX_δ-AVA with Zr-MOLEX_γ-ABA and found that the amino acid compositions of the isolated protein C-terminal peptides were almost the same, but the isolation efficiency was slightly better with Ce-MOLEX_δ-AVA than Zr-MOLEX_γ-ABA (Figure S4).

**Figure 4.**
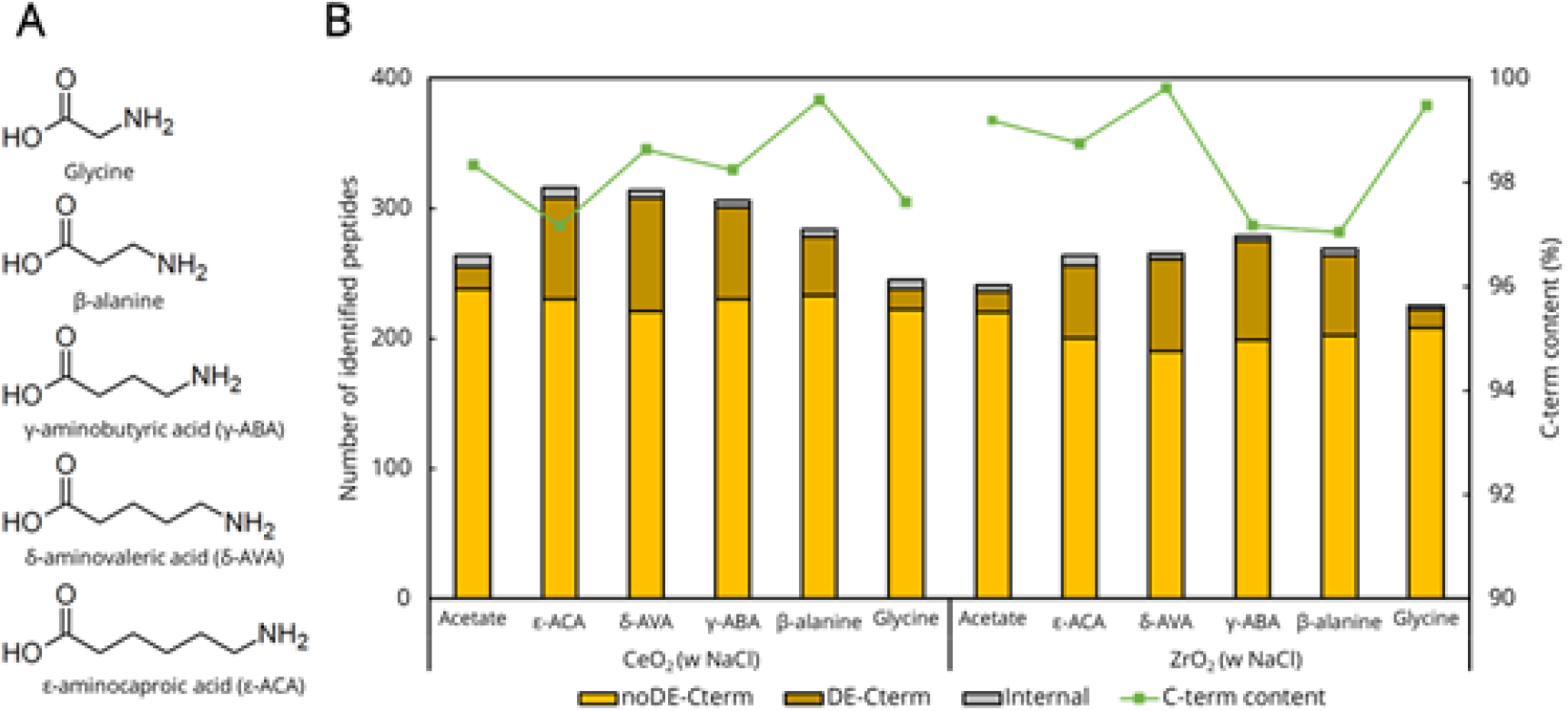
Effect of amino acid additives on the isolation of protein C-terminal peptides by Ce-or Zr-MOLEX chromatography. (A) Amino acids used as mobile phase additives. (B) The identification numbers of non-acidic C-terminal peptides, acidic C-terminal peptides and internal peptides from the flow-through fraction of MOLEX chromatography with different amino acid modifiers. V8-digested HeLa proteins (5 μg) were used for each condition. Protein C-terminal peptide content (%) was calculated as the sum of the peak areas of all identified protein C-terminal peptides divided by that of all identified peptides. The TripleTOF 5600+ system was used to analyze the flow-through pH 5 fractions.

**Figure 5.**
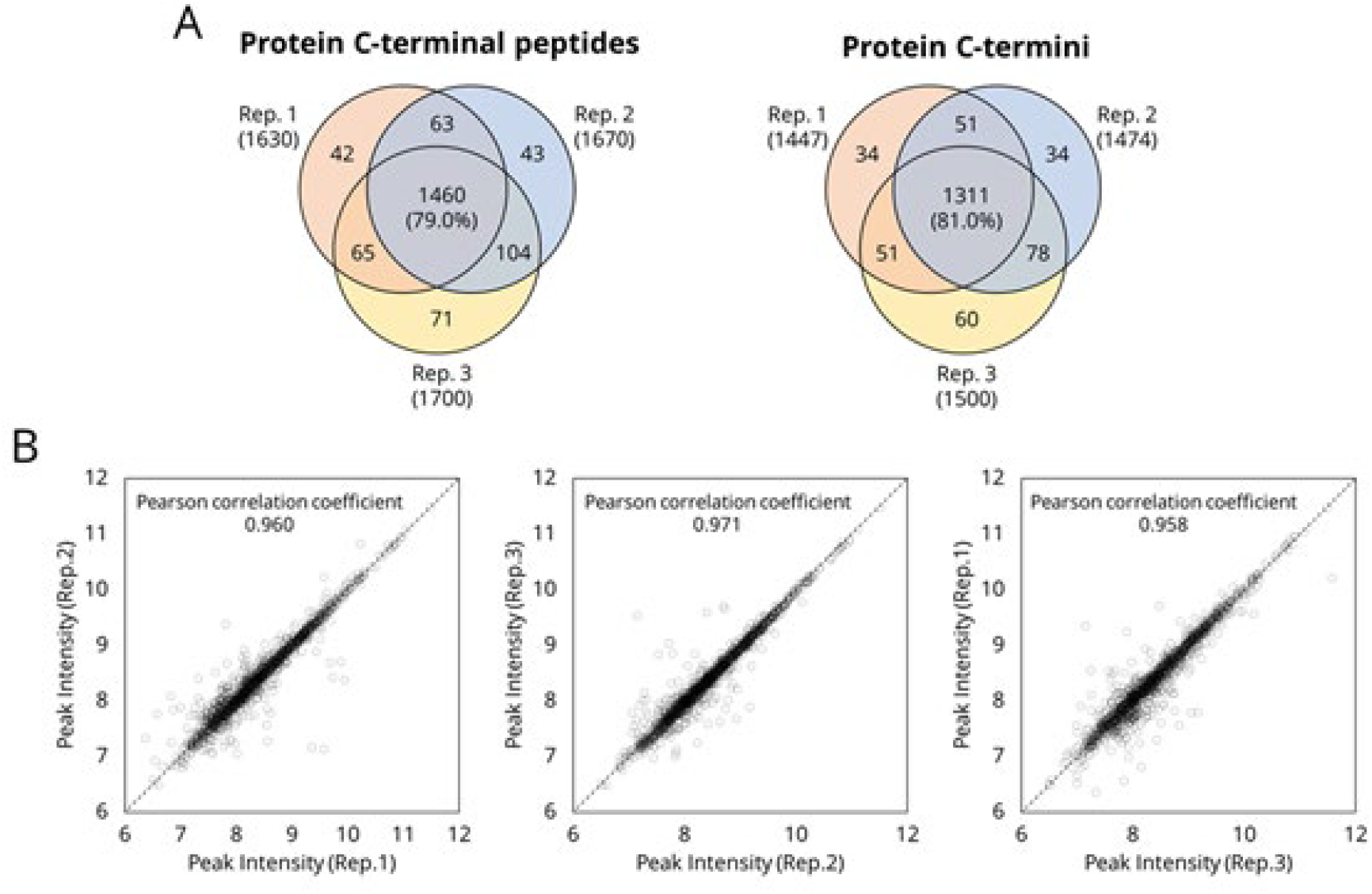
Reproducibility in the Ce-MOLEX isolation step for 10 μg of V8 protease-digested HeLa proteins. (A) The overlaps of the identified protein C-terminal peptide and protein C-termini in triplicate determinations. (B) The scatter plots and the Pearson correlation coefficients for the intensities of the protein C-terminal peptides commonly identified between two measurements. The Orbitrap Exploris 480 system was used.

### Cleavage preference of V8 protease

It is known that the cleavage preference of V8 protease depends upon the digestion buffer,^45,46^ i.e., the protease cleaves the C-terminal side of Glu in ammonium bicarbonate (ABC) buffer, while it cleaves the C-terminal side of Asp in addition to Glu in phosphate buffer.^47^ To check the cleavage selectivity, we prepared three different digestion buffers: 50 mM ABC buffer (pH 7.8), 50 mM Tris-HCl buffer (pH 7.8) and 50 mM phosphate buffer (pH 7.8). HeLa proteins extracted by the PTS buffer containing 12 mM SDC, 12 mM SLS and 100 mM Tris-HCl buffer (pH 7.8) were diluted 5 times with these digestion buffers and incubated overnight with V8 protease (enzyme/substrate ratio = 1:20 (w/w)) at 37 °C. We found that the phosphate buffer showed the best performance in terms of both total identification number and cleavage selectivity, as expected, although the number of peptides with C-terminal Asp was much lower than that of peptides with C-terminal Glu, suggesting that further optimization would be required (Figure S5).

### HeLa C-terminome analysis

Using the HeLa peptides generated by V8 protease in the phosphate buffer as described above, we compared several elution conditions of Ce-MOLEX: (1) 50 μL of aqueous 40% acetonitrile in 600 mM NaCl and 30 mM δ-aminovaleric acid buffers at pH 5, (2) 50 μL x 2 times with the same pH 5 buffer, and (3) 50 μL of the same pH 5 buffer followed by 50 μL of the same buffer but at pH 4. Among them, condition (2) gave a 10% larger number of identified protein C-terminal peptides than condition (1) and a slightly larger number than condition (3) (data not shown). Therefore, we selected condition (2) for the elution of protein C-terminal peptides. Using this condition, we evaluated the isolation reproducibility for 10 μg of V8-digested HeLa proteins in triplicate analyses. A total of 1848 protein C-terminal peptides and 1619 protein C-termini were identified in triplicate (Table S1), and approximately 80% of protein C-terminal peptides as well as protein C-termini were commonly identified (Figure 6A). In addition, we evaluated the peak intensities of the commonly identified protein C-terminal peptides and found that the correlation coefficients were more than 0.95 (Figure 6B), indicating that the reproducibility in isolating protein C-terminal peptides by Ce-MOLEX chromatography was satisfactory.

The obtained results were compared with those of three recently reported C-terminomics studies using HeLa lysates.^23,26,29^ As summarized in Table 1, our method was superior to these reported methods in both sensitivity and efficiency. To deepen the analysis, we used 200 μg of HeLa peptides and isolated the protein C-terminal peptides by Ce-MOLEX followed by high-pH reversed-phase LC fractionation. Subsequent nanoLC/MS/MS analyses of the obtained six fractions resulted in the identification of 2202 protein C-termini with 2.65 % contamination with internal peptides on average, based on the peak intensity (Table S2). To our knowledge, our method provides the largest identification number with the smallest sample size, the smallest contamination of internal peptides, and the shortest isolation time among human protein C-terminomics studies reported to date.

**Table 1.** Comparison of human protein C-terminomics studies.

|  | This study |  | Du et al.<sup>23</sup> | Li et al.<sup>26</sup> |  | Wang et al.<sup>29</sup> |
| --- | --- | --- | --- | --- | --- | --- |
| Sample amount for isolation | HeLa 30 µg | HeLa 200 µg | HeLa 300 µg | HeLa 120 µg | + 293T 120 µg | HeLa 1 mg<sup>a</sup> |
| Required time for isolation | 10 min | 10 min | 300 min | 2880 min |  | 600 min<sup>b</sup> |
| Purity of C-terminal peptide (%) | 93.3 | 97.4 | 16.4 | 22.4 |  | 2.7 - 34 |
| Number of identified protein C-termini | 1619 | 2202 | 781 | 2151 |  | 2162 or 1734<sup>c</sup> |

a) Sample amounts required for propionylation reaction of amino groups.
b) Total time including chemical reactions.
c) After removal of unexpected modifications.

## CONCLUSION

We have developed a one-step isolation method for protein C-terminal peptides based on the CHAMP concept. This C-terminal CHAMP method does not require any chemical modification, and the isolation step takes only 10 minutes using tip-based Ce-MOLEX chromatography. By optimizing the isolation conditions, contamination with internal peptides was minimized, amounting to less than 3%. Although the V8 protease digestion conditions need to be improved to decrease missed cleavages, the current protocol already provides superior results to previously reported methods in terms of the identification number, required sample amount, analysis time, and the purity of the C-terminal peptides. This C-terminal CHAMP method together with our N-terminal CHAMP method^30^ should be useful to investigate biological phenomena regulated by protein terminal modifications.

## ASSOCIATED CONTENT

### Supporting Information

The Supporting Information is available free of charge on the ACS Publications website http://pubs.acs.org

Figure S1, The numbers of identified protein C-terminal and internal peptides using Ce-, Ga-and Al-MOLEX chromatography; Figure S2, Averaged amino acid (%) calculated from the number of each amino acid per peptide divided by the total number of residues for each fraction in Zr-MOLEX chromatography; Figure S3, Dependence of the identification number of protein C-terminal peptides on the sequence position of acidic amino acids (Asp or Glu) in Ce-MOLEX (A) and Zr-MOLEX (B) chromatography with various additives; Figure S4, Comparison of the characteristics of protein C-terminal peptides isolated using Ce-MOLEX_δ-AVA and Zr-MOLEX_γ-ABA; Figure S5, Evaluation of V8 cleavage efficiency in different buffers. (PDF)

Table S1, List of identified protein C-termini in the triplicate samples; Table S2, List of identified protein C-termini in the fractionated samples (XLSX)

## ACKNOWLEDGMENTS

We would like to thank members of the Department of Molecular & Cellular BioAnalysis for fruitful discussions. H. N. was supported by a fellowship for young scientists from the Japan Society for the Promotion of Science (JSPS). This work was supported by the JST Strategic Basic Research Program CREST (No. 18070870), AMED-CREST program (No. JP18gm1010010), JSPS Grants-in-Aid for Scientific Research No. 17H05667, No. 20K21478 and 21H02459 to Y.I and 21J15131 to H.S.

## (TOC)

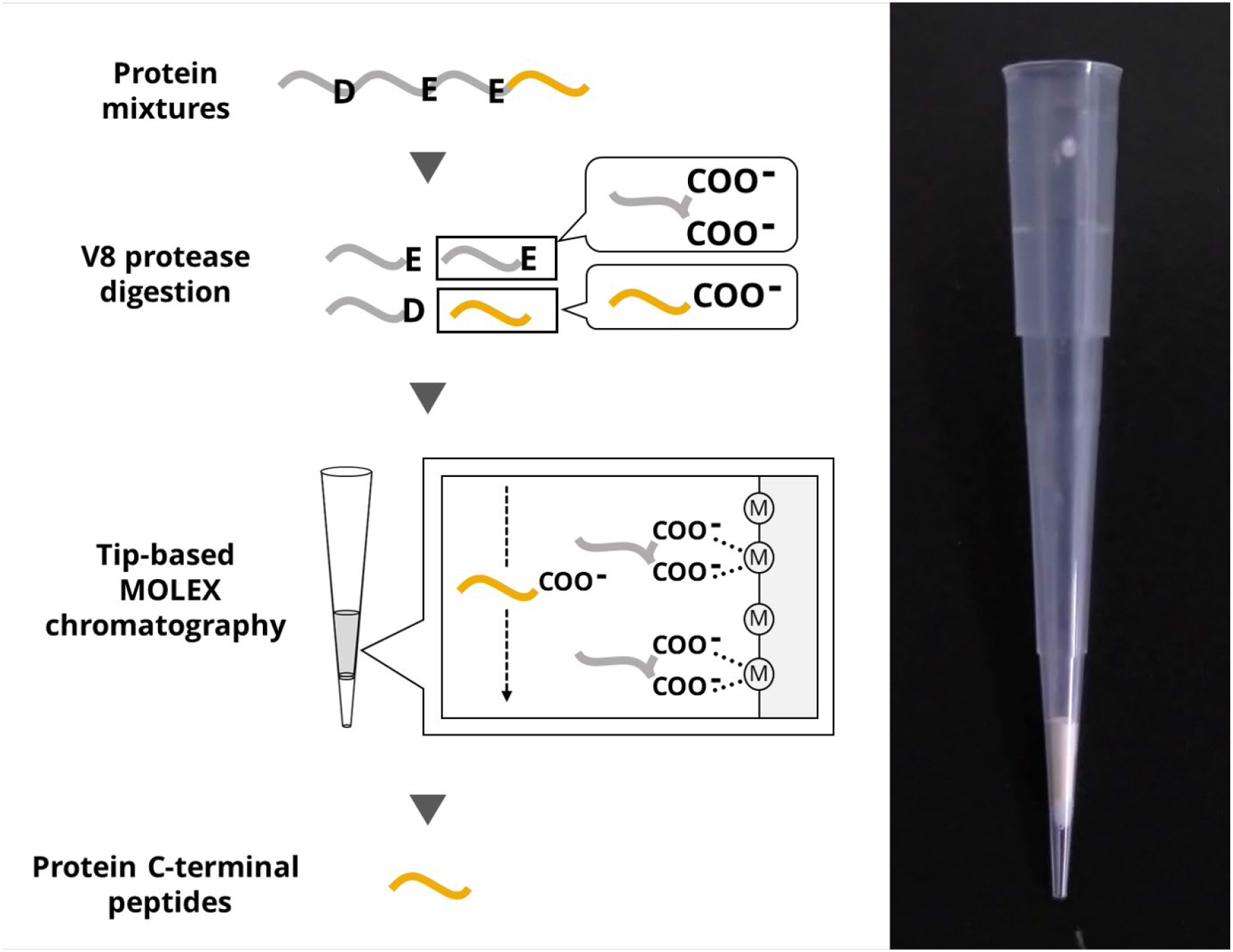

**Figure S1.**
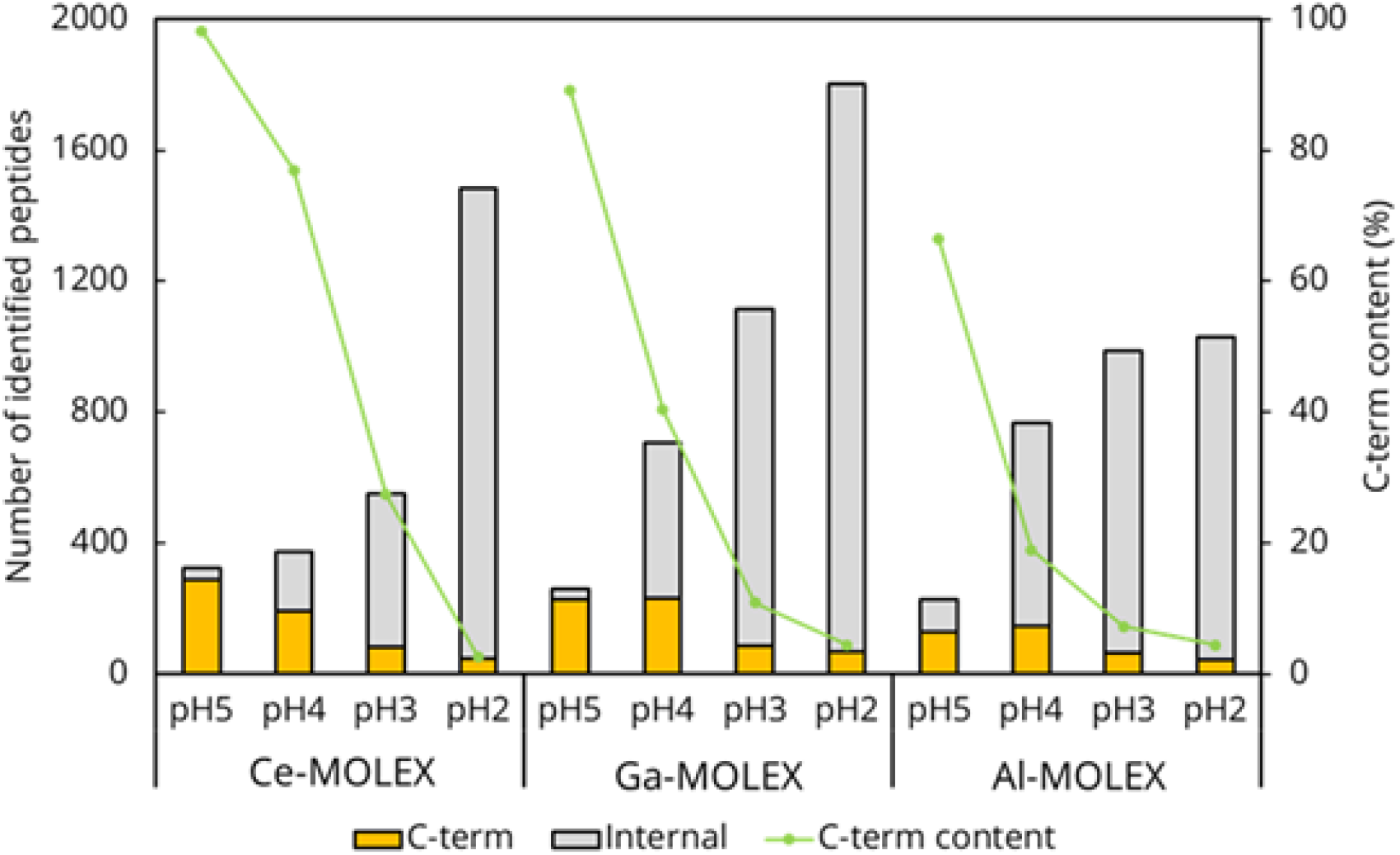
The numbers of identified protein C-terminal and internal peptides using Ce-, Ga- and Al-MOLEX chromatography. Protein C-terminal peptide content (%) was calculated as the sum of the peak areas of all identified protein C-terminal peptides divided by that of all identified peptides. The TripleTOF 5600+ system was used.

**Figure S2.**
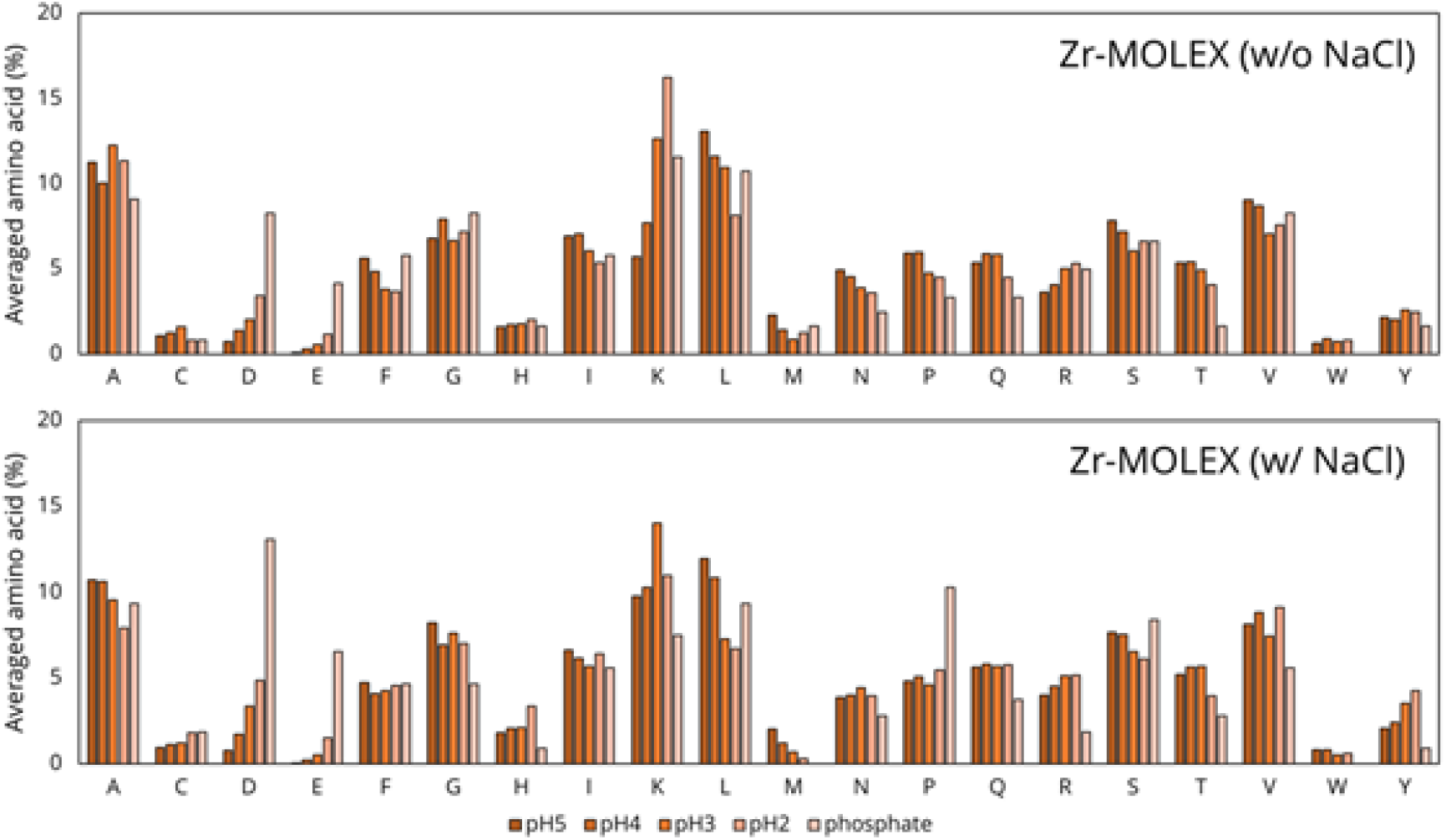
Averaged amino acid (%) calculated from the number of each amino acid per peptide divided by the total number of residues for each fraction in Zr-MOLEX chromatography without NaCl (top) or with NaCl (bottom). The TripleTOF 5600+ system was used.

**Figure S3.**
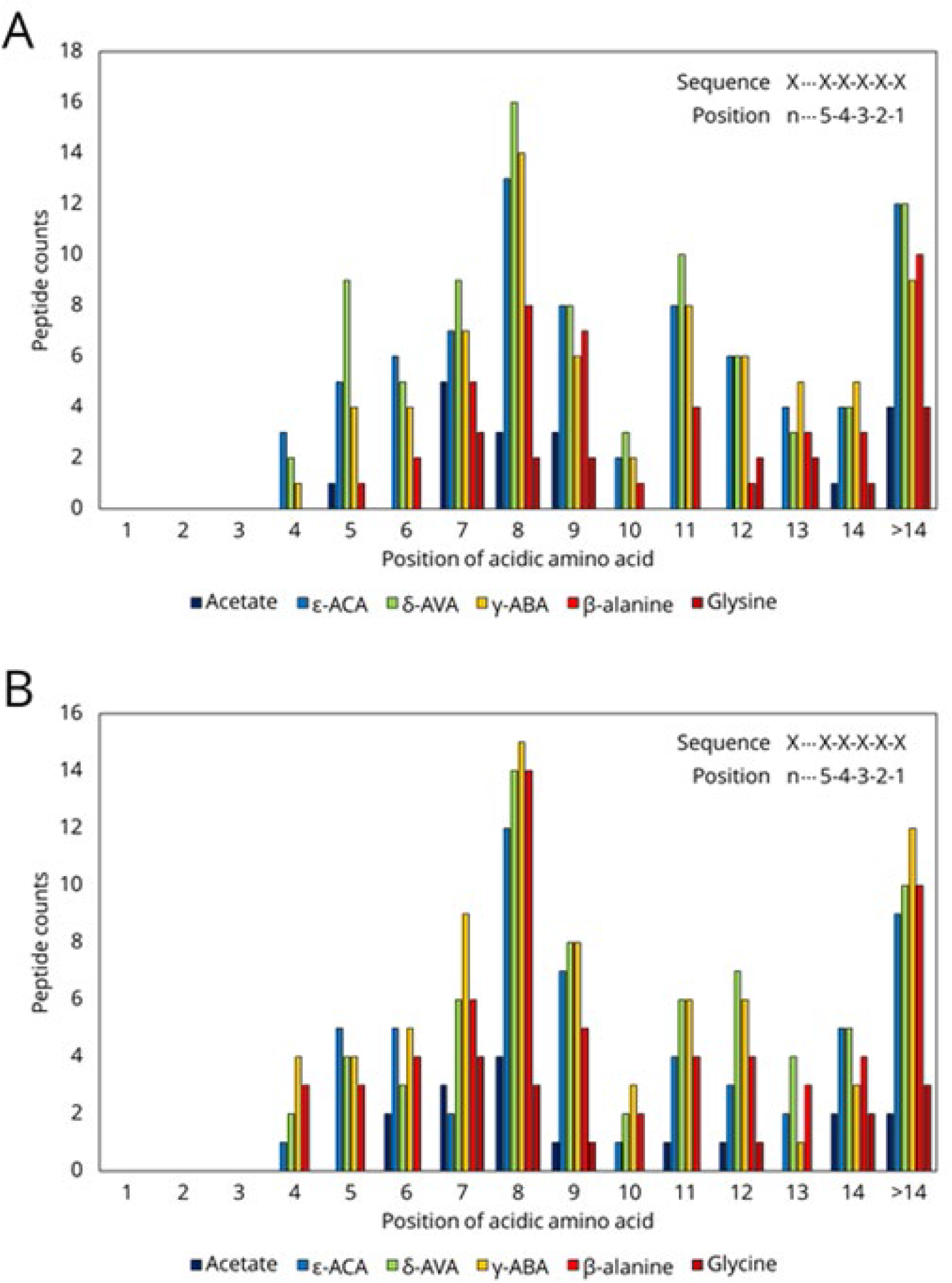
Dependence of the identification number of protein C-terminal peptides on the sequence position of acidic amino acids (Asp or Glu) in Ce-MOLEX (A) and Zr-MOLEX (B) chromatography with various additives. The TripleTOF 5600+ system was used to analyze the flow-through pH 5 fractions.

**Figure S4.**
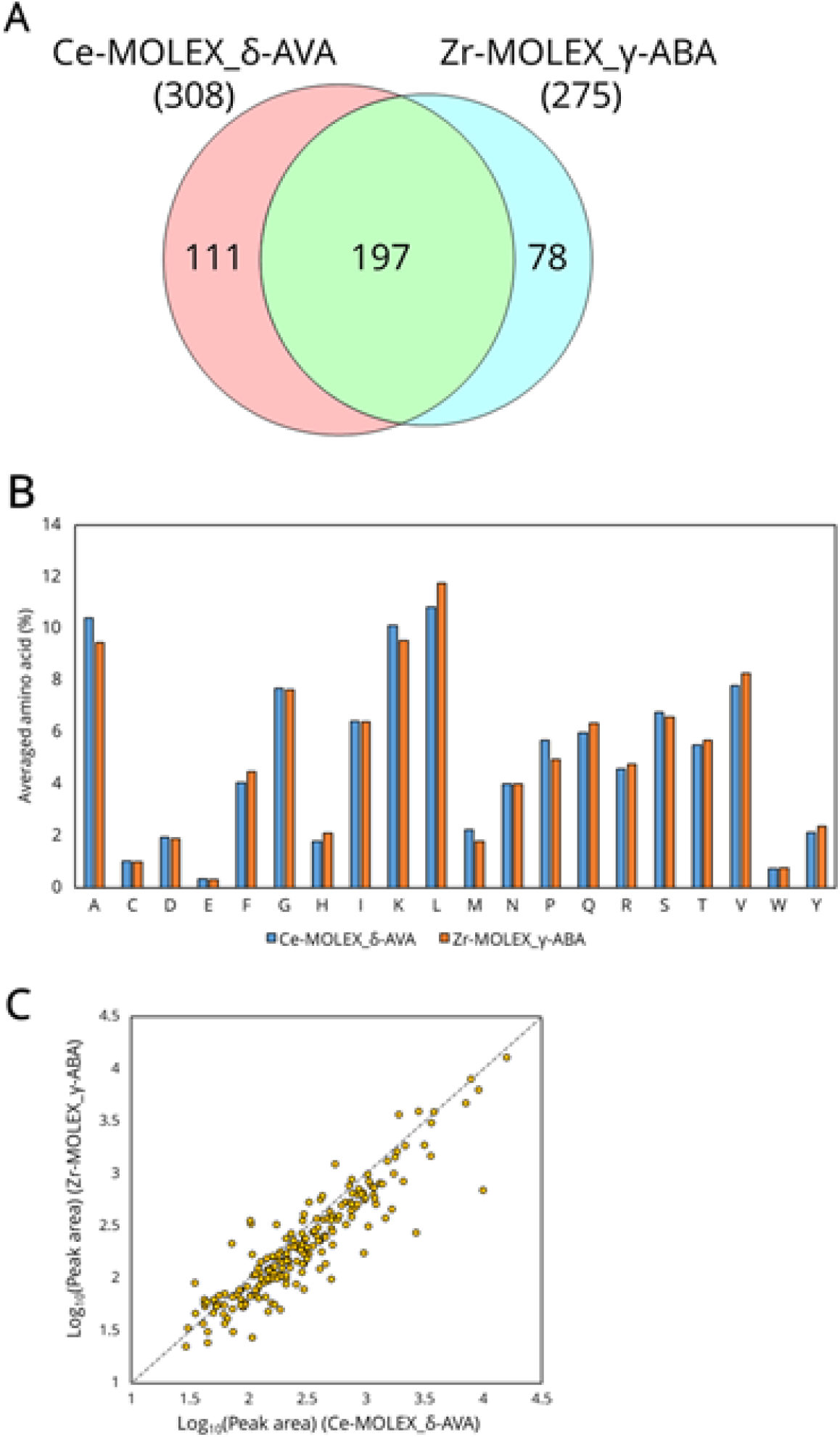
Comparison of the characteristics of protein C-terminal peptides isolated using Ce-MOLEX_δ-AVA and Zr-MOLEX_γ-ABA. (A) Overlap of the identified protein C-terminal peptides. (B) Averaged amino acid (%) calculated from the number of each amino acid per peptide divided by the total number of residues for each fraction in Ce-MOLEX_δ-AVA and Zr-MOLEX_γ-ABA. (C) Comparison of the peak areas of the commonly identified protein C-terminal peptides. The TripleTOF 5600+ system was used to analyze the flow-through pH 5 fractions.

**Figure S5.**
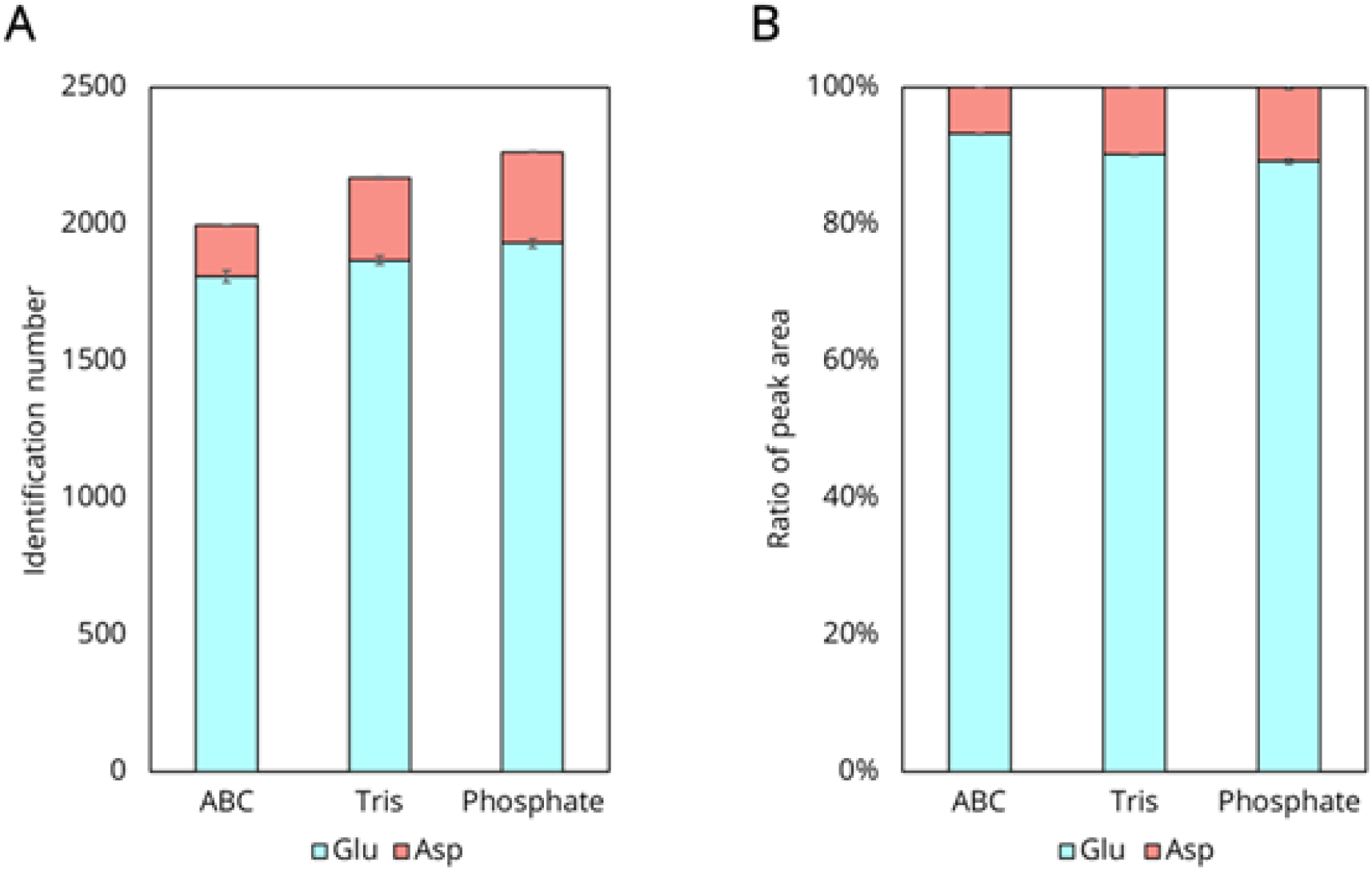
Evaluation of V8 cleavage efficiency in different buffers. (A) The identification numbers of peptides with C-terminal Glu and Asp, (B) the peak area ratio for peptides with C-terminal Glu and Asp. The Q-Exactive system was used.

## REFERENCES

(1) Sriram, S. M.; Kim, B. Y.; Kwon, Y. T. The N-End Rule Pathway: Emerging Functions and Molecular Principles of Substrate Recognition. Nat. Rev. Mol. Cell Biol. 2011, 12 (11), 735–747.

(2) Gawron, D.; Ndah, E.; Gevaert, K.; Van Damme, P. Positional Proteomics Reveals Differences in N-Terminal Proteoform Stability. Mol. Syst. Biol. 2016, 12 (2), 858.

(3) Rogers, L. D.; Overall, C. M. Proteolytic Post-Translational Modification of Proteins: Proteomic Tools and Methodology. Mol. Cell. Proteomics 2013, 12 (12), 3532–3542.

(4) Marino, G.; Eckhard, U.; Overall, C. M. Protein Termini and Their Modifications Revealed by Positional Proteomics. ACS Chem. Biol. 2015, 10 (8), 1754–1764.

(5) Sharma, S.; Schiller, M. R. The Carboxy-Terminus, a Key Regulator of Protein Function. Crit. Rev. Biochem. Mol. Biol. 2019, 54 (2), 85–102.

(6) Chung, J.-J.; Shikano, S.; Hanyu, Y.; Li, M. Functional Diversity of Protein C-Termini: More than Zipcoding? Trends in Cell Biology 2002, 12 (3), 146–150.

(7) Smith, L. M.; Kelleher, N. L.; Consortium for Top Down Proteomics. Proteoform: A Single Term Describing Protein Complexity. Nat. Methods 2013, 10 (3), 186–187.

(8) Smith, L. M.; Thomas, P. M.; Shortreed, M. R.; Schaffer, L. V.; Fellers, R. T.; LeDuc, R. D.; Tucholski, T.; Ge, Y.; Agar, J. N.; Anderson, L. C.; Chamot-Rooke, J.; Gault, J.; Loo, J. A.; Paša-Tolić, L.; Robinson, C. V.; Schlüter, H.; Tsybin, Y. O.; Vilaseca, M.; Vizcaíno, J. A.; Danis, P. O.; Kelleher, N. L. A Five-Level Classification System for Proteoform Identifications. Nat. Methods 2019, 16 (10), 939–940.

(9) Niedermaier, S.; Huesgen, P. F. Positional Proteomics for Identification of Secreted Proteoforms Released by Site-Specific Processing of Membrane Proteins. Biochim. Biophys. Acta: Proteins Proteomics 2019, 1867 (12), 140138.

(10) Tsumagari, K.; Chang, C.-H.; Ishihama, Y. Exploring the Landscape of Ectodomain Shedding by Quantitative Protein Terminomics. iScience 2021, 24 (4), 102259.

(11) Gevaert, K.; Goethals, M.; Martens, L.; Van Damme, J.; Staes, A.; Thomas, G. R.; Vandekerckhove, J. Exploring Proteomes and Analyzing Protein Processing by Mass Spectrometric Identification of Sorted N-Terminal Peptides. Nat. Biotechnol. 2003, 21 (5), 566–569.

(12) Kleifeld, O.; Doucet, A.; auf dem Keller, U.; Prudova, A.; Schilling, O.; Kainthan, R. K.; Starr, A. E.; Foster, L. J.; Kizhakkedathu, J. N.; Overall, C. M. Isotopic Labeling of Terminal Amines in Complex Samples Identifies Protein N-Termini and Protease Cleavage Products. Nat. Biotechnol. 2010, 28 (3), 281–288.

(13) Kaushal, P.; Lee, C. N-Terminomics – Its Past and Recent Advancements. J. Proteomics 2021, 233, 104089.

(14) Tanco, S.; Gevaert, K.; Van Damme, P. C-Terminomics: Targeted Analysis of Natural and Posttranslationally Modified Protein and Peptide C-Termini. Proteomics 2015, 15 (5-6), 903–914.

(15) Dormeyer, W.; Mohammed, S.; van Breukelen, B.; Krijgsveld, J.; Heck, A. J. R. Targeted Analysis of Protein Termini. J. Proteome Res. 2007, 6 (12), 4634–4645.

(16) Xu, G.; Shin, S. B. Y.; Jaffrey, S. R. Chemoenzymatic Labeling of Protein C-Termini for Positive Selection of C-Terminal Peptides. ACS Chem. Biol. 2011, 6 (10), 1015–1020.

(17) Liu, M.; Fang, C.; Pan, X.; Jiang, H.; Zhang, L.; Zhang, L.; Zhang, Y.; Yang, P.; Lu, H. Positive Enrichment of C-Terminal Peptides Using Oxazolone Chemistry and Biotinylation. Anal. Chem. 2015, 87 (19), 9916–9922.

(18) Van Damme, P.; Staes, A.; Bronsoms, S.; Helsens, K.; Colaert, N.; Timmerman, E.; Aviles, F. X.; Vandekerckhove, J.; Gevaert, K. Complementary Positional Proteomics for Screening Substrates of Endo-and Exoproteases. Nat. Methods 2010, 7 (7), 512–515.

(19) Schilling, O.; Barré, O.; Huesgen, P. F.; Overall, C. M. Proteome-Wide Analysis of Protein Carboxy Termini: C Terminomics. Nat. Methods 2010, 7 (7), 508–511.

(20) Nakajima, C.; Kuyama, H.; Tanaka, K. Mass Spectrometry-Based Sequencing of Protein C-Terminal Peptide Using α-Carboxyl Group-Specific Derivatization and COOH Capturing. Anal. Biochem. 2012, 428 (2), 167–172.

(21) Sonomura, K.; Kuyama, H.; Matsuo, E.-I.; Tsunasawa, S.; Nishimura, O. The Specific Isolation of C-Terminal Peptides of Proteins through a Transamination Reaction and Its Advantage for Introducing Functional Groups into the Peptide. Rapid Commun. Mass Spectrom. 2009, 23 (5), 611–618.

(22) Hu, H.; Zhao, W.; Zhu, M.; Zhao, L.; Zhai, L.; Xu, J.-Y.; Liu, P.; Tan, M. LysargiNase and Chemical Derivatization Based Strategy for Facilitating In-Depth Profiling of C-Terminome. Anal. Chem. 2019, 91 (22), 14522–14529.

(23) Du, X.; Fang, C.; Yang, L.; Bao, H.; Zhang, L.; Yan, G.; Lu, H. In-Depth Analysis of C Terminomes Based on LysC Digestion and Site-Selective Dimethylation. Anal. Chem. 2019, 91 (10), 6498–6506.

(24) Kaleja, P.; Helbig, A. O.; Tholey, A. Combination of SCX Fractionation and Charge-Reversal Derivatization Facilitates the Identification of Nontryptic Peptides in C-Terminomics. J. Proteome Res. 2019, 18 (7), 2954–2964.

(25) Helbig, A. O.; Tholey, A. Exopeptidase Assisted N-and C-Terminal Proteome Sequencing. Anal. Chem. 2020, 92 (7), 5023–5032.

(26) Li, Q.; Zhang, Y.; Huang, J.; Wu, Z.; Tang, L.; Huang, L.; Zhang, X. Basic Strong Cation Exchange Chromatography, BaSCX, a Highly Efficient Approach for C-Terminomic Studies Using LysargiNase Digestion. Anal. Chem. 2020, 92 (7), 4742–4748.

(27) Chen, L.; Shan, Y.; Yang, C.; Sui, Z.; Zhang, X.; Zhang, L.; Zhang, Y. Carboxypeptidase B-Assisted Charge-Based Fractional Diagonal Chromatography for Deep Screening of C-Terminome. Anal. Chem. 2020, 92 (12), 8005–8009.

(28) Zhang, Y.; He, Q.; Ye, J.; Li, Y.; Huang, L.; Li, Q.; Huang, J.; Lu, J.; Zhang, X. Systematic Optimization of C-Terminal Amine-Based Isotope Labeling of Substrates Approach for Deep Screening of C-Terminome. Anal. Chem. 2015, 87 (20), 10354–10361.

(29) Wang, Z.; Zhang, L.; Yuan, W.; Zhang, Y.; Lu, H. SAPT, a Fast and Efficient Approach for Simultaneous Profiling of Protein N-and C-Terminome. Anal. Chem. 2021, 93 (30), 10553–10560.

(30) Chang, C.-H.; Chang, H.-Y.; Rappsilber, J.; Ishihama, Y. Isolation of Acetylated and Unmodified Protein N-Terminal Peptides by Strong Cation Exchange Chromatographic Separation of TrypN-Digested Peptides. Mol. Cell. Proteomics 2020, 20, 100003.

(31) Masuda, T.; Tomita, M.; Ishihama, Y. Phase Transfer Surfactant-Aided Trypsin Digestion for Membrane Proteome Analysis. J. Proteome Res. 2008, 7 (2), 731–740.

(32) Rappsilber, J.; Ishihama, Y.; Mann, M. Stop and Go Extraction Tips for Matrix-Assisted Laser Desorption/ionization, Nanoelectrospray, and LC/MS Sample Pretreatment in Proteomics. Anal. Chem. 2003, 75 (3), 663–670.

(33) Rappsilber, J.; Mann, M.; Ishihama, Y. Protocol for Micro-Purification, Enrichment, Pre-Fractionation and Storage of Peptides for Proteomics Using StageTips. Nat. Protoc. 2007, 2 (8), 1896–1906.

(34) Ishihama, Y.; Rappsilber, J.; Andersen, J. S.; Mann, M. Microcolumns with Self-Assembled Particle Frits for Proteomics. J. Chromatogr. A 2002, 979 (1-2), 233–239.

(35) Schilling, B.; Rardin, M. J.; MacLean, B. X.; Zawadzka, A. M.; Frewen, B. E.; Cusack, M. P.; Sorensen, D. J.; Bereman, M. S.; Jing, E.; Wu, C. C.; Verdin, E.; Kahn, C. R.; Maccoss, M. J.; Gibson, B. W. Platform-Independent and Label-Free Quantitation of Proteomic Data Using MS1 Extracted Ion Chromatograms in Skyline: Application to Protein Acetylation and Phosphorylation. Mol. Cell. Proteomics 2012, 11 (5), 202–214.

(36) Kong, A. T.; Leprevost, F. V.; Avtonomov, D. M.; Mellacheruvu, D.; Nesvizhskii, A. I. MSFragger: Ultrafast and Comprehensive Peptide Identification in Mass Spectrometry-Based Proteomics. Nat. Methods 2017, 14 (5), 513–520.

(37) Teo, G. C.; Polasky, D. A.; Yu, F.; Nesvizhskii, A. I. Fast Deisotoping Algorithm and Its Implementation in the MSFragger Search Engine. J. Proteome Res. 2021, 20 (1), 498–505.

(38) da Veiga Leprevost, F.; Haynes, S. E.; Avtonomov, D. M.; Chang, H.-Y.; Shanmugam, A. K.; Mellacheruvu, D.; Kong, A. T.; Nesvizhskii, A. I. Philosopher: A Versatile Toolkit for Shotgun Proteomics Data Analysis. Nat. Methods 2020, 17 (9), 869–870.

(39) Okuda, S.; Watanabe, Y.; Moriya, Y.; Kawano, S.; Yamamoto, T.; Matsumoto, M.; Takami, T.; Kobayashi, D.; Araki, N.; Yoshizawa, A. C.; Tabata, T.; Sugiyama, N.; Goto, S.; Ishihama, Y. jPOSTrepo: An International Standard Data Repository for Proteomes. Nucleic Acids Res. 2017, 45 (D1), D1107–D1111.

(40) Sugiyama, N.; Masuda, T.; Shinoda, K.; Nakamura, A.; Tomita, M.; Ishihama, Y. Phosphopeptide Enrichment by Aliphatic Hydroxy Acid-Modified Metal Oxide Chromatography for Nano-LC-MS/MS in Proteomics Applications. Mol. Cell. Proteomics 2007, 6 (6), 1103–1109.

(41) Sun, S.; Ma, H.; Han, G.; Wu, R. ‘an; Zou, H.; Liu, Y. Efficient Enrichment and Identification of Phosphopeptides by Cerium Oxide Using on-Plate Matrix-Assisted Laser Desorption/ionization Time-of-Flight Mass Spectrometric Analysis. Rapid Commun. Mass Spectrom. 2011, 25 (13), 1862–1868.

(42) Li, Y.; Lin, H.; Deng, C.; Yang, P.; Zhang, X. Highly Selective and Rapid Enrichment of Phosphorylated Peptides Using Gallium Oxide-Coated Magnetic Microspheres for MALDI-TOF-MS and Nano-LC-ESI-MS/MS/MS Analysis. Proteomics 2008, 8 (2), 238–249.

(43) Wolschin, F.; Wienkoop, S.; Weckwerth, W. Enrichment of Phosphorylated Proteins and Peptides from Complex Mixtures Using Metal Oxide/hydroxide Affinity Chromatography (MOAC). Proteomics 2005, 5 (17), 4389–4397.

(44) Tsai, C.-F.; Wang, Y.-T.; Chen, Y.-R.; Lai, C.-Y.; Lin, P.-Y.; Pan, K.-T.; Chen, J.-Y.; Khoo, K.-H.; Chen, Y.-J. Immobilized Metal Affinity Chromatography Revisited: pH/acid Control toward High Selectivity in Phosphoproteomics. J. Proteome Res. 2008, 7 (9), 4058–4069.

(45) Drapeau, G. R.; Boily, Y.; Houmard, J. Purification and Properties of an Extracellular Protease of Staphylococcus Aureus. J. Biol. Chem. 1972, 247 (20), 6720–6726.

(46) Houmard, J.; Drapeau, G. R. Staphylococcal Protease: A Proteolytic Enzyme Specific for Glutamoyl Bonds. Proc. Natl. Acad. Sci. U. S. A. 1972, 69 (12), 3506–3509.

(47) Sørensen, S. B.; Sørensen, T. L.; Breddam, K. Fragmentation of Proteins byS. Aureusstrain V8 Protease. FEBS Letters 1991, 294 (3), 195–197.

